# Impact of PIK3CA gain and PTEN loss on mantle cell lymphoma biology and sensitivity to targeted therapies

**DOI:** 10.1101/2024.04.30.591749

**Authors:** Nardjas Bettazova, Jana Senavova, Kristyna Kupcova, Dana Sovilj, Anezka Rajmonova, Ladislav Andera, Karla Svobodova, Adéla Berkova, Vaclav Herman, Zuzana Zemanova, Lenka Daumova, Alexandra Dolníkova, R. Eric Davis, Marek Trneny, Pavel Klener, Ondrej Havranek

## Abstract

Besides many other mutations in known cancer driver genes, mantle cell lymphoma (MCL) is characterized by recurrent genetic alterations of important regulators of the phosphoinositol-3-kinase (PI3K) cascade including *PIK3CA* gains and *PTEN* losses. To evaluate the biological and functional consequences of these aberrations in MCL, we have introduced transgenic expression of *PIK3CA* (PIK3CA UP) and performed knockout of *PTEN* gene (PTEN KO) in 5 MCL cell lines. The modified cell lines were tested for associated phenotypes including dependence on upstream B-cell receptor (BCR) signaling (by an additional *BCR* knockout). PIK3CA overexpression decreased the dependence of the tested MCL on prosurvival signaling from BCR, decreased levels of oxidative phosphorylation, and increased resistance to 2-deoxy-glucose, a glycolysis inhibitor. Unchanged AKT phosphorylation status and unchanged sensitivity to a battery of PI3K inhibitors suggested that *PIK3CA* gain might impact MCL cells in AKT independent manner. *PTEN* KO was associated with a more distinct phenotype: AKT hyperphosphorylation and overactivation, increased resistance to multiple inhibitors (most of the tested PI3K inhibitors, BTK inhibitor ibrutinib, and BCL2 inhibitor venetoclax), increased glycolytic rates with resistance to 2-deoxy-glucose, and significantly decreased dependence on prosurvival BCR signaling. Our results suggest that the frequent aberrations of the PI3K pathway may rewire associated signaling with lower dependence on BCR signaling, better metabolic and hypoxic adaptation, and targeted therapy resistance in MCL.

**Key point 1:** PIK3CA gain and PTEN loss decrease the dependence of MCL cells on B-Cell Receptor Signaling and anti-apoptotic BCL2.

**Key point 2:** PIK3CA gain and PTEN loss lead to complex metabolic rewiring and increased survival of MCL cells under hypoxia.

## INTRODUCTION

Mantle cell lymphoma (MCL) is a B cell non-Hodgkin lymphoma characterized by a chronically relapsing clinical course(1, 2). Besides the canonical translocation t(11;14)(q13;q32) leading to the overexpression of cyclin D1, many other recurrent molecular-cytogenetic events have been reported in patients with newly diagnosed MCL including lesions of the phosphatidylinositol 3-kinase (PI3K) – protein kinase B (AKT) - mammalian target of rapamycin (mTOR) pathway. These include copy number gains of the P110a catalytic subunit of PI3 kinase (*PIK3CA*), and loss of phosphatase and tensin homologue (*PTEN*)(3-8).

The precise impact of these recurrent aberrations on the biology, survival, and drug resistance of MCL cells remain largely unexplored. Hyperphosphorylation of AKT kinase, because of genetic loss of *PTEN*, has been reported in prognostically adverse MCL blastoid variant(5). Increased expression of *PIK3CA* was associated with MCL progression, while increased PI3K-AKT activity was detectable in patients who failed to respond to ibrutinib, an inhibitor of BTK (Bruton’s tyrosine kinase, a critical kinase involved in B-cell receptor signaling) (9-11). We have recently demonstrated that *PIK3CA* copy number gains represent frequent genetic events associated with MCL relapse after failure of standard immunochemotherapy(12). Because of this pathway deregulation, pharmacological targeting of the PI3K-AKT-mTOR cascade has been repeatedly investigated in MCL, either as a single-agent or in combination with other targeted agents, having promising results (13-20).

In this study, we aimed to investigate the exact consequences of *PIK3CA* gain and *PTEN* loss on the biology of MCL cells and on the sensitivity/resistance to PI3K, BTK, AKT and BCL2 family inhibitors.

## RESULTS

### PIK3CA copy number gain and PTEN deletion are common aberrations in patients with MCL

To determine the prevalence of *PIK3CA* copy number variants, and *PTEN* deletions, we carried out fluorescence *in situ* hybridization (FISH) and array comparative genomic hybridization (aCGH) analyses of 61 consecutive samples obtained from patients with newly diagnosed MCL. Forty-three percent of patients had gain of one copy of *PIK3CA* gene, 7% patients had monoallelic deletion of *PTEN*, one case had both. There was no obvious correlation between *TP53* deletion and *PIK3CA* gain. Interestingly, *PTEN* deletion was found exclusively in the context of concurrent *TP53* deletion, but relatively low numbers of the analyzed patients preclude us to draw any statistically significant correlations The correlation between PTEN deletion and TP53 deletion could not be determined due to the small number of cases with PTEN deletion (Supplemental Table 1).

### PTEN KO but not PIK3CA overexpression show hyperphosphorylation and overactivation of AKT

Overexpression of *PIK3CA* in MINO, Z138, UPF1H, UPF19U, and JEKO-1 *PIK3CA* UP cells, and loss of *PTEN* expression in UPF1H, MINO, Z138, and JEKO-1 *PTEN* KO cells, were confirmed by western blotting. UPF19U *PTEN* Knock-down (*PTEN* KD) manifested a decreased expression rather than complete loss of *PTEN* (Figure 1A, Supplemental Figure 1). Except for JEKO-1, all *PTEN* KO cells had hyperphosphorylation of the key downstream target of PI3K, the AKT kinase (Figure 1A, Supplemental Figure 1). In addition, all *PTEN* KO cells had increased AKT activity by FRET assay (Figure 1B). In contrast, transgenic (over)expression of *PIK3CA* was not associated with consistent changes in AKT phosphorylation or AKT activity (Figure 1C).To determine the extent and potential correlation of *PTEN* expression and AKT phosphorylation in MCL cells, we conducted western blot profiling of 28 primary MCL samples obtained from patients with newly diagnosed and relapsed MCL. Seven out of 28 samples (P3, 6, 15, 16, 19, 20, and 24) had lower expression of *PTEN* and high phosphorylation of AKT (Supplemental Figure 2).

**Figure 1.**
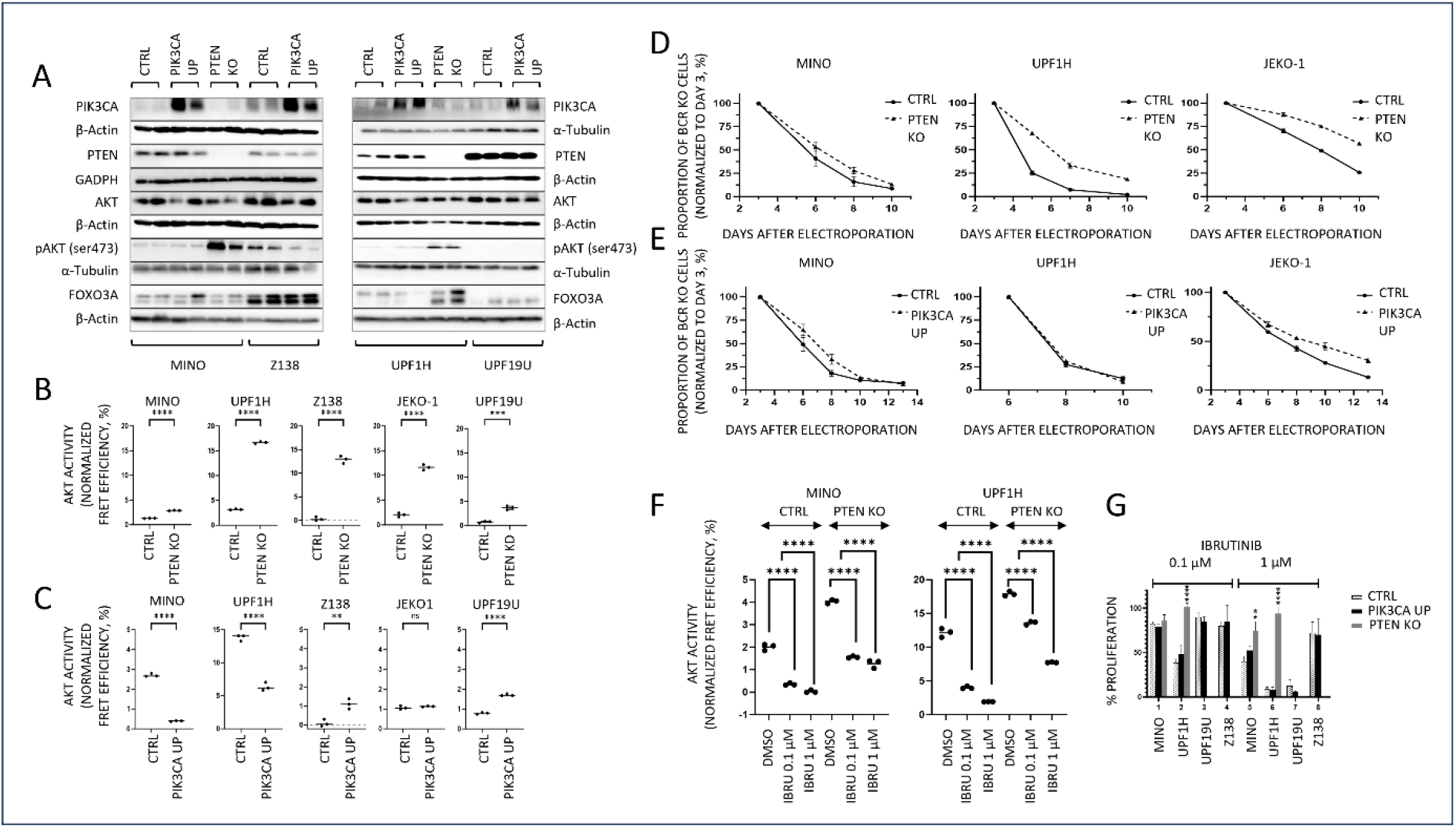
Loss of *PTEN* expression but not *PIK3CA* (over)expression is associated with hyperphosphorylation and overactivation of AKT. On the other hand, both loss of *PTEN* and, to a lesser extent, gain of *PIK3CA* decrease dependance on the pro-survival signaling from BCR. (A) Western blot. Confirmation of the upregulation of *PIK3CA* in the four *PIK3CA* UP clones MINO, Z138, UPF1H, and UPF19U and the loss of *PTEN* in the *PTEN* KO clones MINO and UPF1H. Increase of phospho-AKT (ser 473) expression in both *PTEN* KO clones MINO and UPF1H; for each sample N=2; (B) AKT activity as measured using genetically encoded FRET-based biosensor. Significantly increased AKT kinase activity in all *PTEN* KO clones as compared to respective control cell lines; technical triplicates; (C) FRET assay. Variable effect of *PIK3CA* (over)expression on AKT activity; technical triplicates; (D) *PTEN* loss increases survival of MCL cells with knockout of BCR gene; N=3; (E) transgenic (over)expression of *PIK3CA* in MINO and JEKO-1 but not in UPF1H increases survival of MCL cells with knockout of BCR gene; N=3 (F) AKT activity as measured using genetically encoded FRET-based biosensor. AKT activity in *PTEN* KO clones is higher compared to respective cell lines. In addition, after exposure to BTK inhibitor ibrutinib for 3 hours, AKT activity in *PTEN* KO clones remains higher than AKT activity in the respective CTRL cell lines; N=3; (G) Proliferation assay implemented 72 hours after exposure to Ibrutinib (0.1, 1 μM), the cellular proliferation of the treated cells was normalized to the cellular proliferation of the untreated cells; N=3; data are represented as means ± SD; “ns” means “not significant”, * p<0.05, ** p<0.01, *** p<0.001, **** p<0.0001; (N) represents the number of biological replicates.

### Loss of PTEN expression and gain of PIK3CA both decrease dependence of MCL cells on pro-survival signaling from BCR

We have clearly documented before that BCR signaling critically contributes to PI3K/AKT signaling activation in lymphomas. The degree of PI3K/AKT activation is largely dependent on BCR surface abundance (21). Therefore, we tested if PI3K/AKT hyperactivation (triggered by *PTEN* knockout or *PIK3CA* overexpression) can decrease dependency of MCL cells on the upstream BCR mediated PI3K/AKT activation. We have performed BCR knockout in three *PTEN* KO cell line variants (MINO, UPF1H, and JEKO-1 *PTEN*/BCR KO subclones) and in three *PIK3CA* UP cell line variants (MINO, UPF1H, and JEKO-1 *PIK3CA* UP/BCR KO subclones). Loss of *PTEN* partially decreased dependence on the pro-survival signaling from the BCR (Figure 1D). Partial *PTEN* KO mediated BCR KO rescue was detectable especially in cell lines with large increase of AKT activity following *PTEN* KO (Figure 1B) and was consisted with presumed “chronic active” type of BCR signaling in MCL that activates multiple signaling pathways than just PI3K/AKT (22). Curiously, in two out of three tested *PIK3CA* UP/BCR KO subclones (MINO, and JEKO-1, but not UPF1H), gain of *PIK3CA* also decreased dependence on BCR pro-survival signaling, though to a lower extent in comparison to *PTEN*/BCR KO subclones (Figure 1E). As for the remaining MCL cell lines used in this study, Z138 cells are BCR independent (Supplemental Figure 3) and UPF19U cells do not express BCR.

Importantly, following BTK inhibition by Ibrutinib, *PTEN* KO cells showed lower dependence on BCR-mediated AKT activation in comparison to parental cell lines (as detected by genetically encoded FRET-based AKT activity biosensor, Figure 1F, supplemental figure 4)(23). This implies that loss of *PTEN* overcomes Ibrutinib mediated decrease of AKT activity which should translate into BTK inhibition resistance. Indeed, proliferation assays confirmed that *PTEN* KO cells were more resistant to ibrutinib (Figure1G). On the other hand, *PIK3CA* overexpression did not consistently increase AKT activity and caused a paradoxical AKT activity decrease in several cell lines (Figure 1C). In accordance with this, AKT activity decrease following ibrutinib treatment was not rescued by *PIK3CA* overexpression (Supplemental Figure 5) and no changes in the sensitivity to Ibrutinib were detected (Figure 1G).

### Loss of PTEN and gain of PIK3CA affect metabolic activity and survival of MCL cells under hypoxia

The PI3K-AKT signaling pathway plays a pivotal role in regulation of cell energy-metabolic pathways. Therefore, we used Seahorse analyzer to assess potential impact of *PIK3CA* gain or *PTEN* loss on metabolic reprogramming of MCL. In UPF1H cells, both *PTEN* KO and *PIK3CA* overexpression decreased basal as well as maximal respiration (Figure 2A). Interestingly, these two genetic alterations had different effects on glycolysis in UPF1H cells. *PTEN* KO massively increased basal glycolysis (with almost no further glycolytic reserve left). Basal glycolysis was not affected by *PIK3CA* overexpression, however, *PIK3CA* overexpression decreased overall glycolytic capacity and glycolytic reserve (Figure 2B). In MINO cells, *PIK3CA* overexpression had a similar effect as in UPF1H (decreased maximal respiration and decreased glycolytic parameters), however it was much less pronounced. PTEN KO did not affect respiration or glycolysis in UPF1H cells. (Figures 2C-D). Substantial decrease of mitochondrial function, accompanied by increased basal glycolysis in *PTEN* KO cells, suggest increased glycolysis to ox/phos ratio in *PTEN* KO as well as *PIK3CA* UP UPF1H cells, which was not clearly detectable in MINO cells.

**Figure 2.**
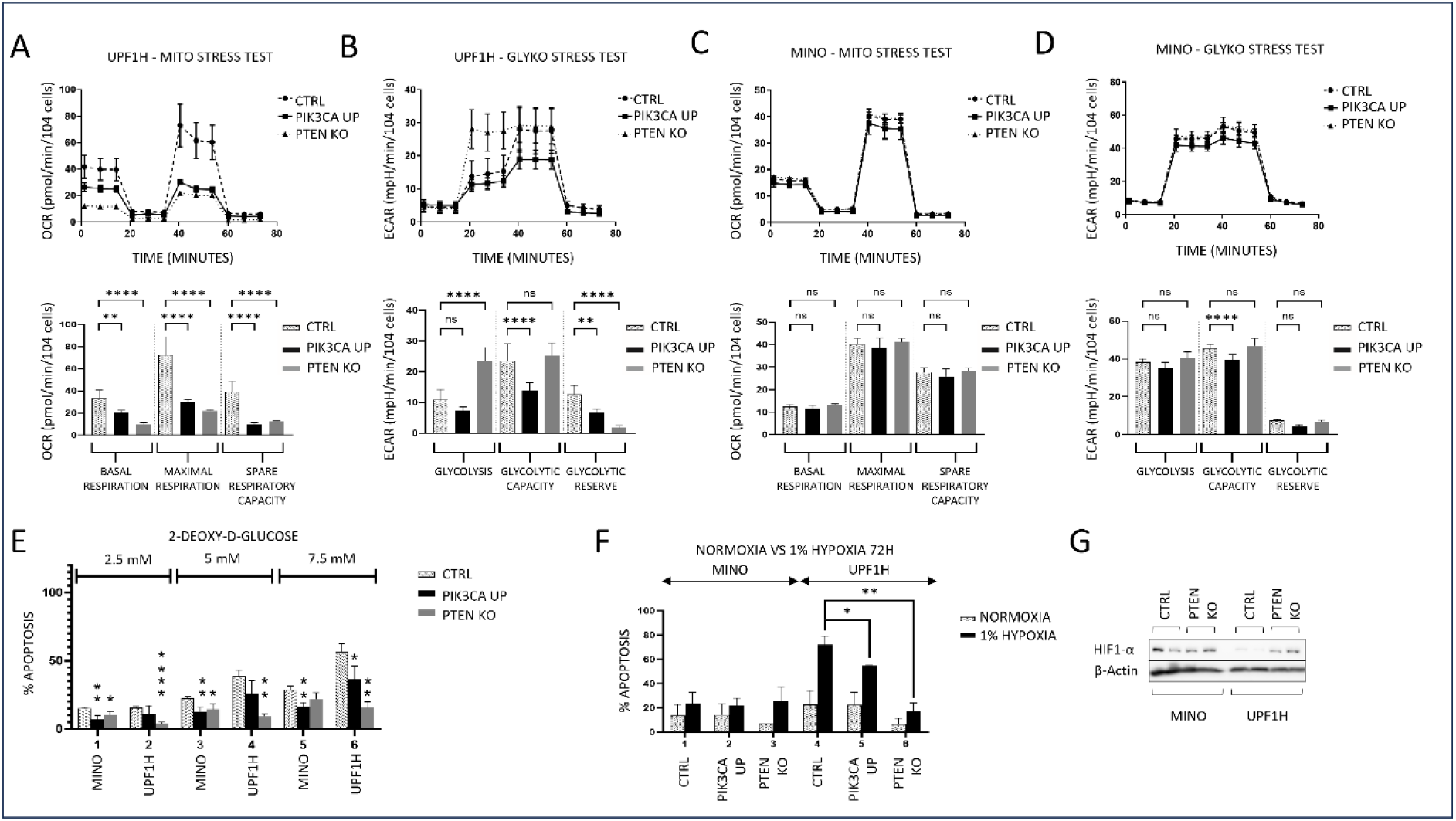
*PTEN* KO and *PIK3CA* UP clones have changed the activity of key energy-metabolic pathways and have increased survival under hypoxia. (A-D) Mito stress tests and glycol stress tests performed using SeaHorse analyzer; (A) Decrease in the basal respiration, maximal respiration, and spare respiratory capacity in UPF1H *PIK3CA* UP and *PTEN* KO cells as compared to UPF1H unmodified cell line; measured by oxygen consumption rate (OCR); (B) Increase in basal glycolysis in UPF1H *PTEN* KO cells, decrease of glycolytic capacity in UPF1H *PIK3CA* UP cells, and decrease of glycolytic reserve in both tested variants; measured by extracellular acidification rate (ECAR); (C) No significant changes of mitochondrial function in in MINO *PIK3CA* UP modified cells compared to MINO CTRL; (D) Slight decrease of glycolysis parameters in in MINO *PIK3CA* UP modified cells compared to MINO CTRL; (E) Number of apoptotic cells 24 hours after exposure to the inhibitor of glycolysis 2-Deoxy-D-Glucose (2-DG) (2.5, 5, 7.5 mM); apoptosis of the treated cells was normalized to the apoptosis of the untreated cells; N=3; (F) Increased survival of *PTEN* KO and *PIK3CA* UP clones after 72 h exposure to 1% hypoxia; apoptosis of the cells cultured for 72 hours under 1% hypoxia (as well as apoptosis of the cells cultured for 72 hours in parallel under normoxia) was normalized to the apoptosis of the cells before placement into hypoxia; N=3; (G) Western blot analysis of HIF1-alpha in PTEN clones KO compared to respective cell lines; for each sample N=2; data are represented by means ± SD; “ns” means “not significant”, * p<0.05, ** p<0.01, *** p<0.001, **** p<0.0001; (N) represents the number of biological replicates.

Functionally, both *PTEN* KO and *PIK3CA* UP UPF1H and MINO cell lines were generally more resistant to 2-deoxy-glucose (2-DG), an inhibitor of glycolysis (Figure 2E). Like metabolic changes, this phenotype was more prominent in UPF1H cells.

Increased glycolytic rate is characteristic for malignant cells growing under hypoxia. Therefore, we tested survival of modified cell lines under 3-day exposure to hypoxia (1% oxygen). UPF1H *PTEN* KO and *PIK3CA* UP cells were both more resistant to long-term exposure to 1% hypoxia (Figure 2F). UPF1H *PTEN* KO cells, but not UPF1H *PIK3CA* UP cells had higher levels of HIF1-alpha protein by western blotting as compared to the parental UPF1H cells (Figure 2G). In contrast, MINO cells were generally resistant to low tensions of oxygen, and, therefore, no significant differences were observed between *PTEN* KO or *PIK3CA* UP modified cells and the parental MINO cells regarding differences in growth in hypoxia and HIF1-alpha upregulation (Figure 2F-G).

### MCL PIK3CA UP and PTEN KO clones have changed sensitivity to BH3 mimetics

Giving the important anti-apoptotic role of the PI3K/AKT pathway and the clinical potential of BH3 mimetics, we also tested the cytotoxic efficacy of selected BH3 mimetics in the context of *PTEN* knockout and *PIK3CA* overexpression.

While all tested *PTEN* KO cells were more resistant to the BCL2 inhibitor venetoclax (VEN), their sensitivity to the BCL-XL inhibitor A1155463 was increased (Figure3A-B). Two out of four tested *PIK3CA* UP cells (UPF1H and MINO) were more resistant to VEN and S63845, a MCL1 inhibitor (Figure 3A,3C).

**Figure 3.**
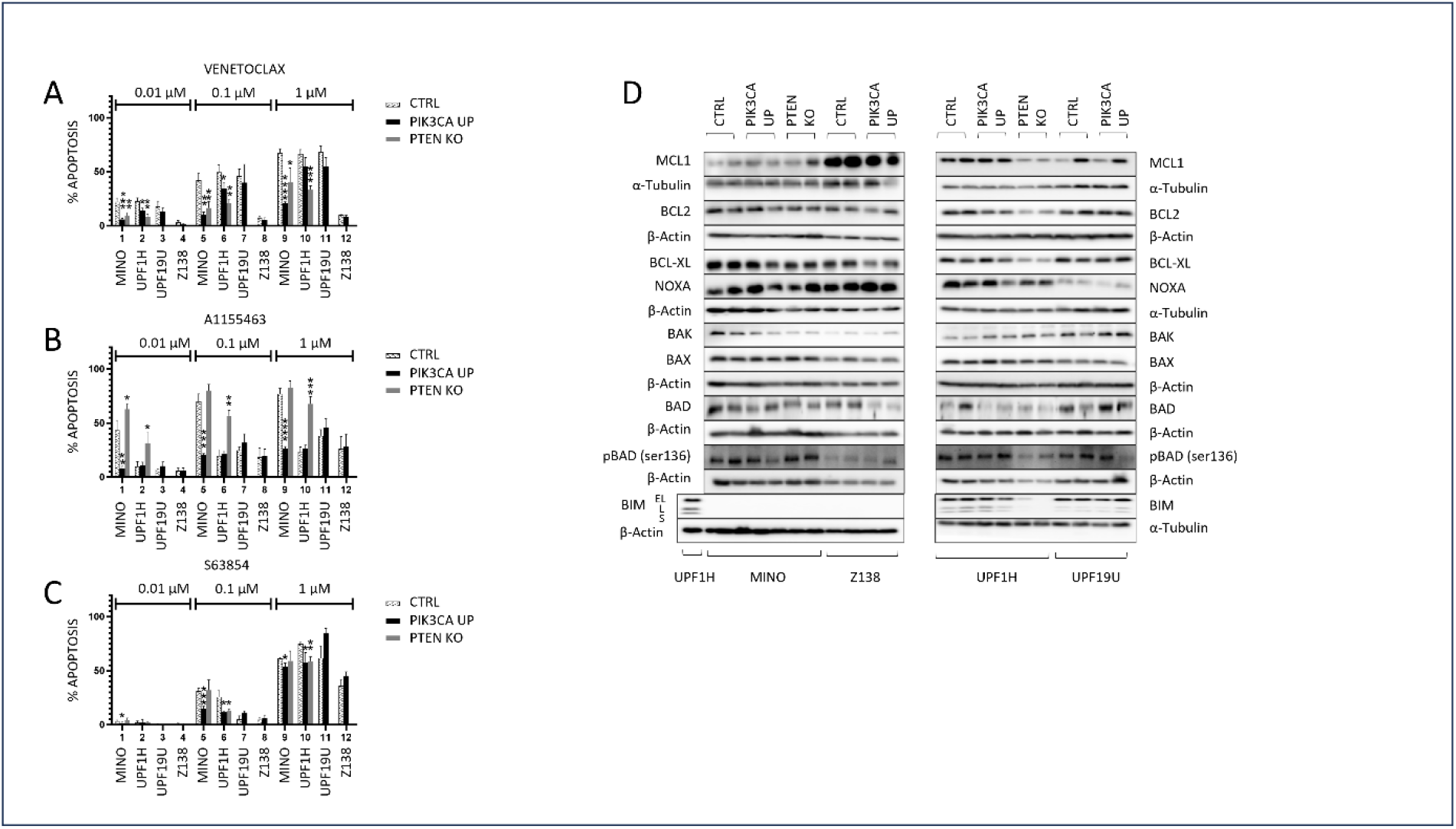
*PIK3CA* UP and *PTEN* KO leads to different sensitivities to BH3 mimetics (A-C) Number of apoptotic cells 24 hours after exposure to the BH3 mimetics venetoclax (0.01, 0.1, 1 μM), S63845 (0.01, 0.1, 1 μM), and A1155463 (0.01, 0.1, 1 μM). Apoptosis of the treated cells was normalized to the apoptosis of the untreated cells. N=3; (D) Western blot analysis of selected BCL2 family proteins in *PIK3CA* and *PTEN* clones compared to the respective MCL cell lines; for each sample N=2; data are represented by means ± SD; * p<0.05, ** p<0.01, *** p<0.001, **** p<0.0001; (N) represents the number of biological replicates.

To decipher the molecular mechanisms underlying the observed changes in the sensitivity to the tested BH3 mimetics, we implemented western blot analysis to determine the expression of key BCL2 family proteins, and immunoprecipitated BCL2 and BCL-XL to identify their respective binding partners.

UPF1H *PTEN* KO cells had significantly decreased expression of proapoptotic BIM (MINO cells have biallelic deletion of BIM). MINO *PTEN* KO cells expressed more phospho-BAD (Figure 3D). Immunoprecipitation experiments demonstrated decreased amounts of all analyzed proapoptotic BCL2 proteins on BCL-XL. We observed decreased interactions of BIM, BAD, BID, BAK, and BAX with BCL-XL in the UPF1H *PTEN* KO cells. In the MINO PTEN KO cells, only BAX showed decreased interaction with BCL-XL in comparison with the parental line (Supplemental Figure 6).

### PTEN KO and to a lesser degree PIK3CA overexpression mediated resistance to a panel of PI3K inhibitors

To explore the therapeutic consequences of *PTEN* loss and *PIK3CA* gain, we implemented a cytotoxic screen of the established *PIK3CA* UP and *PTEN* KO cells using a panel of selected pathway inhibitors including PI3K inhibitors (idelalisib, duvelisib, AZD8186, AZD8835, alpelisib, and copanlisib), and an AKT inhibitor capivasertib. For the functional assays, we used two *PTEN* KO (UPF1H, MINO) and four *PIK3CA* UP cell lines variants (UPF1H, MINO, UPF19U, and Z138). Loss of *PTEN* correlated with increased resistance to PI3K inhibitors (Figure 4 A-F). While UPF1H *PTEN* KO cells were more resistant to all tested PI3K inhibitors, MINO *PTEN* KO cells were more resistant to AZD8186, and copanlisib. *PTEN* KO clones of both lines retained sensitivity to AKT inhibitor capivasertib (Figure 4G). With few exceptions, gain of *PIK3CA* was not associated with significant changes in the sensitivity to PI3K inhibitors or capivasertib.

**Figure 4.**
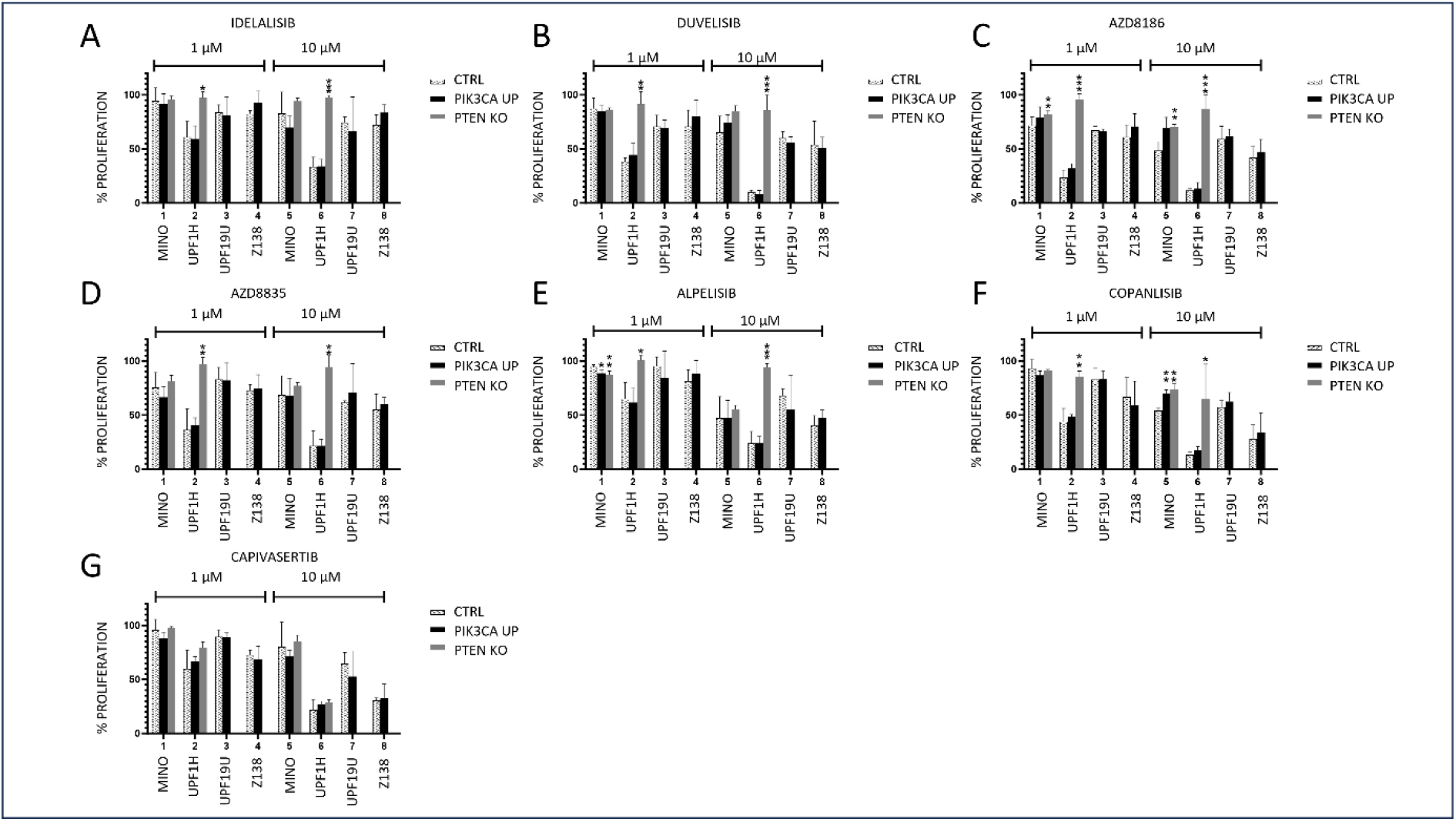
*PTEN* KO and to a lesser degree *PIK3CA* overexpression mediated resistance to a panel of PI3K inhibitors (A-G) Proliferation assays implemented 72 hours after exposure to the indicated agents: (A) Idelalisib (1, 10 μM), (B) Duvelisib (1, 10 μM), (C) AZD8186 (1, 10 μM), (D) AZD8835 (1, 10 μM), (E) Alpelisib (1, 10 μM), (F) Copanlisib (0.01, 0.1 μM), (G) Capivasertib (1, 10 μM); the cellular proliferation of the treated cells was normalized to the cellular proliferation of the untreated cells; N=3; data are represented as means ± SD; * p<0.05, ** p<0.01, *** p<0.001, **** p<0.0001; (N) represents the number of biological replicates.

## DISCUSSION

Aberrant overactivation of the PI3K-AKT signaling pathway has been repeatedly reported in MCL and was associated with blastoid morphology and resistance to BTK inhibitors(5, 7, 9, 10, 24, 25). In this study, we confirmed that gain of *PIK3CA* and deletion of *PTEN* are recurrent aberrations in patients with newly diagnosed MCL. Interestingly, the most frequent type of aberration was a gain of just one copy of *PIK3CA* gene found in 43% of patients. We may speculate that fine-tuning of the PI3K-AKT pathway, rather than massive uncontrolled overactivation, is needed for accurate benefits from its activation in MCL cells. This is in line with previous reports by Shojaee et al. who demonstrated that both the blockage and massive overactivation of PI3K-AKT is detrimental for malignant cells(26). Importantly, FISH analysis was performed only in patients with bone marrow involvement (≥ 20% infiltration). Therefore, frequency of *PIK3CA* gains in patients with low or no bone marrow infiltration remains unknown. Frequencies of *PIK3CA* gains and *PTEN* losses at the first or subsequent disease relapses also remain elusive. However, our recent whole exome sequencing-based study on clonal evolution of 25 MCL patients at diagnosis and at first relapse (after failure of standard immunochemotherapy) revealed that *PIK3CA* gain is one of the most frequently identified copy number variant associated with MCL relapse after failure of standard immunochemotherapy(12).

Surprisingly, transgenic (over)expression of *PIK3CA* was not associated with measurable changes in the phosphorylation or activity levels of AKT kinase or increased resistance to a wide spectrum of tested PI3K inhibitors or the BTK inhibitor ibrutinib in the majority of tested MCL cell lines. On the other hand, most of the tested MCL cell lines with *PIK3CA* overexpression were less sensitive to BH3 mimetics. Molecular mechanisms underlying observed increase in resistance to BH3 mimetics, however, remain elusive. Importantly, *PIK3CA* gain variably decreased dependence of the tested MCL cell lines on pro-survival signaling from the BCR, decreased levels of ox/phos, and increased resistance to 2-DG. Generally, these changes may suggest that *PIK3CA* gain contributes to increased survival of MCL cells, especially under hypoxia. Even though precise molecular mechanisms that mediate these observations remain largely elusive, unchanged activity and phosphorylation status of AKT suggest that consequences of *PIK3CA* gain are mediated independently of AKT. This is supported by a study in endometrial carcinoma. Frederik Holst *et al*. showed that endometrial tumors with *PIK3CA* amplification were associated with aggressive behavior, however, had decreased levels of activated AKT at the same time(27). AKT independent cancer promoting effect was reported also for *PIK3CA* mutations in breast cancer(28).

In contrast to *PIK3CA* (over)expression, knockout of *PTEN* was associated with a distinct phenotype including hyperphosphorylation and overactivation of AKT kinase, increased resistance to most of the tested PI3K inhibitors, resistance to the BTK inhibitor ibrutinib and BCL2 inhibitor venetoclax, markedly decreased dependence on the pro-survival BCR signaling, and metabolic reprogramming of MCL cells with increased glycolytic rate and increased resistance to 2-DG. Importantly, sensitivity to AKT inhibitor capivasertib remained unchanged, which is consistent with data published thus far(29). Interestingly, sensitivity to BCL-XL inhibitor A1155463 was increased. The observations from the western blot analyses (i.e., downregulation of BCL2 and BIM proteins in UPF1H PTEN KO, and downregulation of BAK in MINO PTEN KO), and immunoprecipitation experiments (i.e., decreased interaction of BIM with BCL2 in UPF1H *PTEN* KO) may at least partially explain increased resistance of *PTEN KO* clones to venetoclax. However, molecular mechanisms underlying the observed increased sensitivity to BCL-XL inhibition remain elusive.

Even though *PTEN* deletion is rather a rare event in newly diagnosed MCL, frequency of this aberration in relapsed patients remains unknown. Importantly, it has been shown that also non-genetic mechanisms at the transcriptional and post transcriptional level may result in decreased activity of PTEN phosphatase in cancer (30, 31). Indeed, previous studies focusing on PTEN expression in MCL lymphoma showed mainly reduction of its expression in majority of cases in contrast to a complete PTEN loss (5, 7, 8). These results were confirmed in our own study where PTEN loss was not frequent, however, multiple primary MCL samples showed substantial decrease of its expression.

Importantly, our data show that loss of *PTEN* may negatively correlate with sensitivity to BTK inhibitors. However, this correlation must be confirmed on primary MCL samples before and after failure of BTKi therapy.

We proved that both aberrations (i.e., gain of *PIK3CA*, and loss of *PTEN*) might lead to metabolic reprogramming of MCL cells, which results in significant changes in the rates of glycolysis and ox/phos, increased resistance to 2-DG, and increased survival under hypoxia. Previous studies demonstrated that hypoxia adaptation was associated with allelic loss or inactivation of *PTEN*(32, 33). Conversely, loss of *PTEN* facilitated hypoxia induced factor 1 (HIF1)-mediated gene expression(34-36). Metabolic reprogramming belongs to hallmarks of cancer(37-39). Overactivation of PI3K pathway promotes glycolysis in an AKT dependent and independent manner. One example of AKT independent activation of glycolysis is the effect of the above-mentioned *PIK3CA* mutation(40, 41). Together with the observed decreased dependence on the pro-survival signaling from BCR, we propose that both *PTEN* loss and *PIK3CA* gain foster survival of MCL cells under hypoxia.

## CONCLUSION

Our results suggest that frequent aberrations of the PI3K pathway observed in MCL may rewire associated signaling with lower dependence on BCR signaling, better metabolic and hypoxic adaptation, and targeted therapy resistance.

## MATERIALS AND METHODS

### Cell lines and clones

MCL cell lines (MINO, Z138, JEKO-1) were purchased from Deutche Sammlung of Microorganisms and Zellkulturen (DSMZ), or American Tissue Culture Collection (ATCC), UPF1H and UPF19U were derived in our laboratory from leukemized peripheral blood of patients with chemotherapy-resistant MCL. Generation of cells with transgenic *PIK3CA* (over)expression (*PIK3CA* UP) or with *PTEN* knockout (*PTEN* KO) and cultivation conditions are further described in the supplemental material.

### Cytotoxic agents

Specific PI3K inhibitors, AKT inhibitor, BTK inhibitor and BCL2 family inhibitors are described in the supplemental material.

### Cytogenomic analyses

Copy number alteration analyses of the *PIK3CA, PTEN* and *TP53* genes were performed either by interphase fluorescence *in situ* hybridization (I-FISH) or by array-comparative genomic hybridization/single-nucleotide polymorphism (aCGH/SNP), depending on the availability of material, as part of the routine diagnostic procedure. Detailed protocol description is provided in the supplemental material.

### Generation of cell lines with target genes knockout

Detailed protocol description is provided in the supplemental material.

### Generation of cell lines with target genes overexpression

Detailed protocol description is provided in the supplemental material.

### Cell lines transfection

To introduce all plasmids, we used electroporation according to the manufacturer’s instructions (Neon, ThermoFisher Scientific). 1.2 million cells per sample were used for the 100 μl electroporation system. Cells were washed once by PBS and resuspended in 120 μl of electroporation R buffer together with plasmid DNA (12 μg for single KO plasmid for GFP and BCR KO; 5 + 5 + 5 μg for three plasmids necessary for *PTEN* KO GFP KI; and 6 + 4 μg for sleeping beauty donor and transposase plasmids, respectively, for stable overexpression) to perform electroporation. Cells were then resuspended in 3 ml of appropriate cell culture media without antibiotics for further experiments or selection. Electroporation conditions for individual cell lines were as follows (voltage, pulse length, number of pulses): Jeko-1 1500V, 20ms, 1 pulse; Mino 1300V, 30ms, 1 pulse; UPF1H 1500V, 10ms, 3 pulses; UPF19U 1500V, 10ms, 3 pulses; Z138 1600V, 20ms, 1 pulse.

### Measurement of AKT activity in living cells

Detailed protocol description is provided in the supplemental material.

### Apoptosis and proliferation assays

Detailed protocol description is provided in the supplemental material.

### Western blotting

Detailed protocol description is provided in the supplemental material.

### Coimmunoprecipitation assay

Detailed protocol description is provided in the supplemental material.

### In vitro assays under hypoxia

The in vitro experiments under hypoxic conditions were implemented in Coy hypoxic box. Detailed protocol description is provided in the supplemental material.

### Oxygen Consumption Rate (OCR) and Extracellular Acidification Rate (ECAR) assays

OCR and ECAR measurements were performed using a Seahorse XFe96 analyzer (Agilent Technologies). On the day of the experiment, cells were seeded on XFe96 cell culture microplates coated with Cell-Tak (Corning, 5×10^4^ cells/well), in Seahorse base RPMI medium and incubated at 37°C, without CO_2_ for 1hr. The base medium (180μl/well) was supplemented with 500mM pyruvate, 2mM L-glutamine, 10mM glucose for OCR measurement, and with 2mM L-glutamine for ECAR measurement. The mito stress test and glycolytic stress test are further described in the supplemental material.

### Statistical analysis

To assess the statistical significance of experiments, unpaired student’s t-test or ordinary one-way ANOVA were used (GraphPad Prism-10). p-value < 0.05 represents statistical significance. Densitometry was carried out using Image-Lab Bio-Rad-6.1 software.

## Author Contribution

PK and OH designed the research, analyzed data, and supervised preparation of the manuscript, NB performed most of the research, and wrote the paper, JS, KK, and AR derived genetically modified clones, LA, DS implemented Seahorse analyses, KS, AB, and ZZ did FISH analyses, LD and AD did hypoxic experiments, MT, and RED analyzed data and reviewed the manuscript.

