## Supplemental data files for "Impact of PIK3CA gain and PTEN loss on mantle cell lymphoma biology and sensitivity to targeted therapies"

**SUPPLEMENTAL MATERIALS AND METHODS**

*Cell lines and patient samples*

MINO, Z138, UPF1H, UPF19U, JEKO-1 cell lines with transgenic *PIK3CA* (over)expression (*PIK3CA* UP) or with *PTEN* knock-out (*PTEN* KO) were derived as described below. Cells were maintained in Iscove's modified Dulbecco's medium (IMDM) supplemented with 15% fetal bovine serum (FBS) and 1% penicillin/streptomycin. Primary MCL cells were obtained from leukemized blood, infiltrated lymph nodes, malignant ascites, pleural effusion, and infiltrated bone marrow of 28 patients with so far untreated or relapsed / refractory (R/R) MCL. Approval was obtained from all patients according to the (WMA) Declaration of Helsinki. The project was approved by the institutional ethics committee of the General University Hospital in Prague under number 60/20.

*Cytotoxic agents*

Specific PI3K inhibitors including idelalisib (PI3Kδ inhibitor), duvelisib (PI3K/γ δ inhibitor), AZD8835 (PI3K α/δ inhibitor), AZD8186 (PI3Kβ/δ inhibitor), alpelisib (PI3Kα inhibitor), copanlisib (pan-PI3K inhibitor), and others; capivasertib (pan-AKT inhibitor), ibrutinib (BTK inhibitor), venetoclax (BCL2 inhibitor), S63854 (MCL1 inhibitor), and A1155463 (BCL-XL inhibitor) were purchased from MedChemExpress. 2-Deoxy-D-Glucose (2-DG, inhibitor of glycolysis) was provided by SIGMA.

*Cytogenomic analyses*

I-FISH analyses were performed with commercially available DNA probes SPEC PIK3CA/CEN 3 DC (**ZytoVision GmbH, Bremerhaven, Germany**), Vysis LSI PTEN/CEP 10 and Vysis LSI TP53 (17p13.1)/CEP 17 (Abbott Molecular, Des Plaines, IL, USA) according to the manufacturers' protocols. At least 200 interphase nuclei were analyzed by two independent observers. The cut off level for positive values were determined on samples obtained from 10 cytogenetically normal persons and were found to be 5% (mean ± 3SD) for losses (deletions, monosomies), and 2.5% (mean ± 3SD) for gains (trisomies, amplifications). aCGH/SNP analyses were performed with a SurePrint G3 Cancer CGH+SNP Microarray, 4x180K (Agilent Technologies, Santa Clara, USA) . The QIAamp DNA Blood Mini Kit (Qiagen Inc., Hilden, Germany) was used to isolate genomic DNA from bone-marrow cells stored in fixative. The concentration and quality of the isolated DNA were confirmed with a NanoDrop 2000 spectrophotometer (Thermo Fisher Scientific, Waltham, MA). The array slides were scanned with a microarray scanner system (G2565CA, Agilent Technologies) and analyzed with Agilent Cytogenomics v5.2.0.20 software (Agilent Technologies).

*Generation of cell lines with target genes knock out*

*PTEN* KO was generated using CRISPR/Cas9 system exactly as we did before(1). Briefly, we combined double strand DNA cut at the *PTEN* translation initiation site and provided repair template plasmid for homologous repair (HR) based insertion of GFP-STOP-pA DNA sequence. In result, GFP is expressed instead of *PTEN* and serves as a marker for viable sorting of *PTEN* KO cells. To introduce double strand DNA break at the *PTEN* translation initiation site, we used a "paired nickases" approach(2). *PTEN* target sequences (PTEN_1 TTGACCTGTATCCATTTCTGCGG and PTEN_2 TTGATGATGGCTGTCATGTCTGG) were cloned into a chimeric plasmids pX335-U6-Chimeric_BB-CBh-hSpCas9n(D10A), coding for Cas9 D10A as well as gRNA (Addgene plasmid #42335)(3) and co-electroporated (see below) them with the above-mentioned HR repair template plasmid(1). If necessary for consequent AKT activity measurement (see below), GFP was knocked out using similar CRISP/Cas9 system with single GFP target sequence (GGGCGAGGAGCTGTTCACCGGGG) cloned into a pX330-U6-Chimeric_BB-CBh-hSpCas9 (Addgene plasmid # 42230, coding for Cas9 as well as gRNA)(3). GFP negative *PTEN* KO cells were then sorted to purity for consequent analyses(4). BCR KO cells were generated as we did previously(1). We used gRNAs targeting the constant region of immunoglobulin heavy (IgH) chain of cell line specific BCR isotype. We used the above mentioned px330 plasmid with the following IgH M target sequence for all tested cell lines: AGATGAGCTTGGACTTGCGGGGG. The BCR KO cell growth and the change of their proportion in cell culture was measured starting three days after electroporation using flow cytometry (staining the BCR with anti-human IgM FITC antibody, ThermoFisher Scientific #H15001).

*Generation of cell lines with target genes overexpression*

Similarly as we did before, we have used the sleeping beauty transposon system(5) for stable overexpression of *PIK3CA* and to express the AKT activity reporter (see below) in selected cell lines(1, 6). Briefly, the WT PIK3CA cDNA was cloned into donor plasmid pSBbi Pur (Addgene plasmid #60523)(5) and AKT activity reporter was cloned into pSBbi-Pur (Addgene plasmid #60523)(5) or pSBbi-Neo (Addgene plasmid #60525)(5) donor palasmids. These donor plasmids (6 µg) were co-electroporated with 4 µg of transposase coding plasmid pCMV(CAT)T7-SB100 (Addgene plasmid #34879)(7). After three days of culture without antibiotics, selection with appropriate antibiotics was initiated with the following concentrations: puromycin at final concentration of 2 µg/mL and geneticin at final concentration of 200 µg/mL.

*Measurement of AKT activity in living cells*

To measure AKT activity in live cells, we used a genetically encoded Förster resonance energy transfer (FRET) biosensor as we developed and used before(1, 4). The AKT activity reporter (Lyn-Akt AR2-EV, Addgene plasmid #125199) has the following structure: cell membrane targeting sequence (from Lyn)

- mCerulean3 - FHA1 phospho-amino acid binding domain – EV flexible linker – part of FOXO domain (naturally phosphorylated by AKT) - cpVenus[E172]. AKT mediated phosphorylation of its FOXO part leads to its binding to FHA1 domain, change in reporter confirmation, and increase in FRET. As we published previously, we have used flow cytometry for FRET measurement in living cells using in house R package (fRet) to calculate the absolute FRET efficiency (E) values reflecting the AKT activity(1, 4). FRET measurement and calculations were performed as per the fRet package manual using cells expressing appropriate calibration controls: mCerulean3 only, cpVenus[E172] only, FRET high control, and FRET low control (to calculate necessary set up dependent coefficients for each individually performed experiment)(4).

To compare cells with *PTEN* KO or *PIK3CA* UP to their parental counterparts, five hundred thousand cells per sample were resuspended in fresh media, incubated one hour in regular cell culture incubator (37°C with 5% of $CO_2$), and FRET measured for each sample immediately after removing them from the incubator using flow cytometry (Cytoflex, Beckma Coulter). To measure the AKT activity after ibrutinib treatment, five hundred thousand cells per each sample were pre-incubated (37°C with 5% of $CO_2$) for 3 hours with the ibrutinib concentration of 1 µM and 0.1 µM in total amount of 3ml cell culture cell media directly in flow cytometry tubes and FRET measured for each sample immediately after removal from the incubator using flow cytometry (Cytoflex, Beckma Coulter).

*Apoptosis and proliferation assays*

Apoptosis was measured by standard Annexin-V / propidium iodide (PI) assay. Briefly, on day 1 the cells were resuspended into 96-well plate (100 000 cells per well). Each drug was added in 3 different concentrations. After 24 hours of incubation, Annexin V FITC (Exbio) and PI (Sigma) were added to detect apoptotic and necrotic cells. The measurements were carried out by flow cytometry (BD FACS CANTO II). The percentage of necrotic and apoptotic cells was calculated as previously reported(6).

Proliferation was measured by commercially available WST-8 based cell proliferation assay according to the manufacturer´s recommendations. Briefly, on day 1, the cells were resuspended into 96-well plate (50 000 – 100 000 cells per well). The tested PI3K and AKT inhibitors were added at concentrations 1 and 10 µM (Copanlisib at 0,01 and 0,1 µM), BTK inhibitor at 0,1 and 1 µM and BCL2 family inhibitors at 0,01, 0,1 and 1 µM. After 72 hours of incubation, WST-8 reagent from Quick Cell Proliferation Assay Kit (BioVision) was added, for a further 3 hours incubation. Absorbance of samples was measured on ELISA reader. The proliferation curve was calculated as previously reported(8).

*Western blotting*

Samples were lysed in Ripa buffer (150 mM NaCl, 0,1 % SDS,1mM EDTA,0,5% Sodium Deoxycholate, 1% Triton X-100, pH 7.4). A protease (Sigma) and a phosphatase inhibitor (Roche) were added. Protein concentration was determined using Pierce BCA Protein Assay (ThermoFisher scientific) .20 µg of sample was mixed with laemelli sample buffer (BioRAD) containing mercaptoethanol and boiled for 5 min. Duplicate samples were separated on 10%,12% and 15% SDS-PAGE gels. After electrophoresis, proteins were blotted onto 0.22 µM PVDF membranes (advansta). Membranes were incubated for 1 h in 1xPBS containing 0.1% Tween-20 and 5% non-fat dried milk. Samples were incubated with primary antibodies overnight then with secondary antibodies for 30 min. To detect bands WesternBright ECL HRP substrate (advansta) was used. The membranes were imaged by ChemiDoc™MP Imaging system (BioRAD). The following antibodies were used in our study: PIK3CA (# 4249), PTEN (# 5384), phospho-PTEN ser 380 (# 9551), total AKT (# 9272), phospho-AKT ser 473 (# 4060), FOXO3A (# 12829), phospho-FOXO3A ser 253 (# 9466), GSK3-β (# 12566), phospho-GSK3-β ser 9 (# 9336), BIM (# 2933), BAK (# 12105), BAX (# 2774), BCL-XL (# 2764), MCL1 (# 39224) , phospho-BAD ser 136 (# 9295) and HIF1α (# 14179)were from Cell Signaling technology. B-Actin (# 6276), c-MYC (# 32072), and α-Tubulin (# 7291) from ABCAM. BCL2 (# 610539) from BD Transduction Laboratories.BCL2 (#783) from SantaCruiz. GAPDH (# G8795) from SIGMA. BAD polyclonal (# PA5-11403) from Thermo-Fisher Scientific. NOXA polyclonal (# 2437) from ProSci.

*Coimmunoprecipitation assay*

The immunoprecipitation was performed according to the previously mentioned protocol(9). To determine the interaction between both proteins, Anti-BIM antibody was used to target the immunoprecipitated anti-Bcl-2 antibody (# 783) (SantaCruiz). Anti-BAD, Anti-BIM, anti-BID (# 2002) (cell signaling), anti-BAK and anti BAX were used to target immunoprecipitated anti-BCL-XL antibody (# 32370) (ABCAM)

*In vitro assays under hypoxia*

On day 1, 45 x $10^4$ cells from each cell line resuspended in 3 ml medium were pipetted into two separate 6-well plates. One plate was incubated under normoxic conditions ($O_2$ concentration=20%) and one plate under hypoxic conditions ($O_2$ concentration=1%). After 72 h incubation samples were taken from both plates and an apoptosis assay was carried out (for more go to paragraph apoptosis assay)

*Oxygen Consumption Rate (OCR) and Extracellular Acidification Rate (ECAR) assays*

The mito stress test was performed by series of injections, starting with oligomycin (1 µM, ATP synthase inhibitor), followed by CCCP (2 µM, mitochondrial uncoupler), and a combination of antimycin and rotenone (0.5 µM, inhibitors of RCIII and RCI). The glycolytic stress test was run by series of injections, starting with glucose (10mM, substrate of glycolysis), followed by oligomycin (1 µM, inhibits ATP synthase), and 2-deoxy-D-glucose (50 mM, glycolysis inhibitor). OCR/ECAR were measured after each injection. After the assay, the cells were stained with Hoechst and counted using Cytation5 instrument. OCR/ECAR values were normalized to the same cell number.

**SUPPLEMENTAL RESULTS**

**Supplemental table 1.** FISH/aCGH of 61 primary MCL patients with > 20% bone marrow infiltration, 43% have *PIK3CA* gain and 7% have *PTEN* monoallelic loss. N/A stands for "not analyzed".

| Patient | FISH or aCGH | *PIK3CA* | *PTEN* | *TP53* |
|---|---|---|---|---|
| 1 | aCGH | Gain | N | DEL |
| 2 | aCGH | N | N | DEL |
| 3 | aCGH | N | DEL | DEL |
| 4 | aCGH | N | N | DEL |
| 5 | aCGH | Gain | N | N |
| 6 | aCGH | N | N | DEL |
| 7 | aCGH | N | N | N |
| 8 | aCGH | N | N | DEL |
| 9 | aCGH | N | N | N |
| 10 | aCGH | Gain | N | DEL |
| 11 | aCGH | Gain | N | DEL |
| 12 | aCGH | N | N | DEL |
| 13 | aCGH | N | N | DEL |
| 14 | aCGH | N | N | N |
| 15 | aCGH | Gain | N | DEL |
| 16 | aCGH | Gain | DEL | N/A |
| 17 | aCGH | N | N | N |
| 18 | aCGH | N | N | DEL |
| 19 | aCGH | Gain | N | DEL |
| 20 | aCGH | N | N | N |
| 21 | aCGH | Gain | N | N |
| 22 | aCGH | Gain | N | DEL |
| 23 | aCGH | Gain | N | N |
| 24 | aCGH | Gain | N | N |
| 25 | aCGH | N | N | N |
| 26 | aCGH | Gain | N | N |
| 27 | aCGH | Gain | N | DEL |
| 28 | aCGH | N | N | N |
| 29 | aCGH | N | N | N |
| 30 | aCGH | Gain | N | N |
| 31 | aCGH | Gain | N | N |
| 32 | aCGH | Gain | N | N |
| 33 | aCGH | Gain | N | N |
| 34 | aCGH | N | N | N |
| 35 | aCGH | Gain | N | DEL |
| 36 | FISH | Gain | N | N |
| 37 | FISH | N | N | DEL |
| 38 | FISH | N | N | N |
| 39 | FISH | N | N | N |
| 40 | FISH | Gain | N | N |
| 41 | FISH | N | N | N |
| 42 | FISH | N | N | N |
| 43 | FISH | Gain | N | N |
| 44 | FISH | N | N | N |
| 45 | FISH | N | N | N |
| 46 | FISH | N | DEL | DEL |
| 47 | FISH | Gain | N | N |
| 48 | FISH | Gain | N | DEL |
| 49 | FISH | N | N | N |
| 50 | FISH | Gain | N | N |
| 51 | FISH | N | N | N |
| 52 | FISH | Gain | N | DEL |
| 53 | FISH | Gain | Gain | N |
| 54 | FISH | N | N | N |
| 55 | FISH | N | N | N |
| 56 | FISH | N | DEL | DEL |
| 57 | FISH | N | N | DEL |
| 58 | FISH | N | N | N |
| 59 | FISH | N | N | N |
| 60 | FISH | N | N | N |
| 61 | FISH | N | N | DEL |

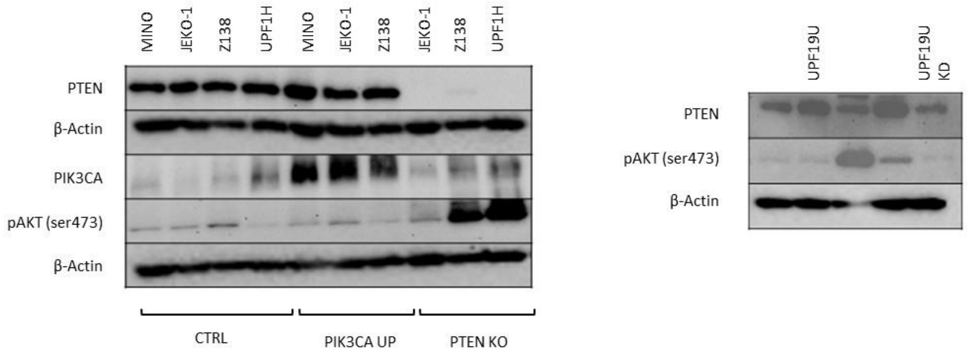


**Supplemental figure 1.** Western blot showing loss of *PTEN* and hyperphosphorylation of AKT in JEKO-1 and Z138 *PTEN* KO cells and overexpression of *PIK3CA* in JEKO-1 *PIK3CA* UP cells. Decreased expression of *PTEN* in UPF19U *PTEN* KD with no changes in the phosphorylation of AKT.

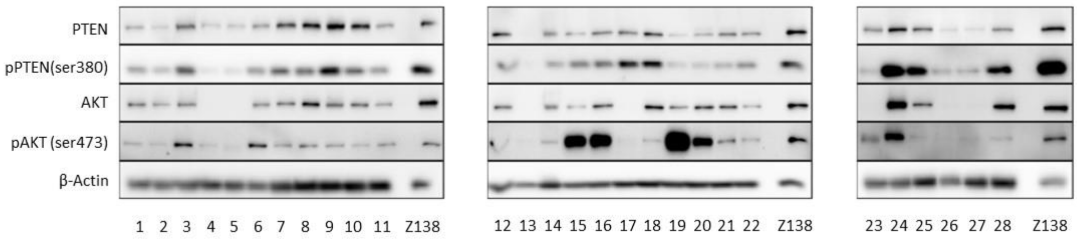


**Supplemental figure 2.** Western blot analysis of 28 primary MCL samples. Detection of PTEN, pPTEN (ser380), AKT and pAKT (ser473) expressions. Samples P3, 6, 15, 16, 19, 20, and 24 have lower expression of *PTEN* and high phosphorylation of AKT.

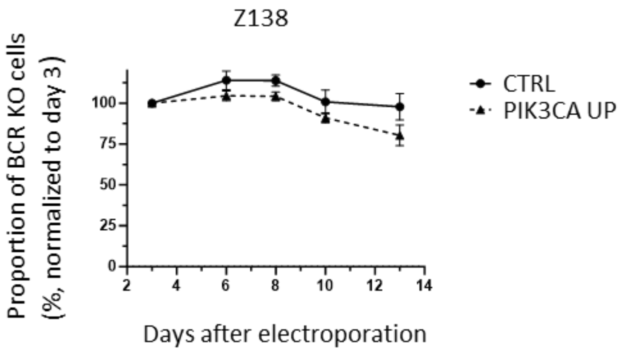


**Supplemental figure 3.** Knock out of BCR in Z138 cell line demonstrates that Z138 is BCR independent.

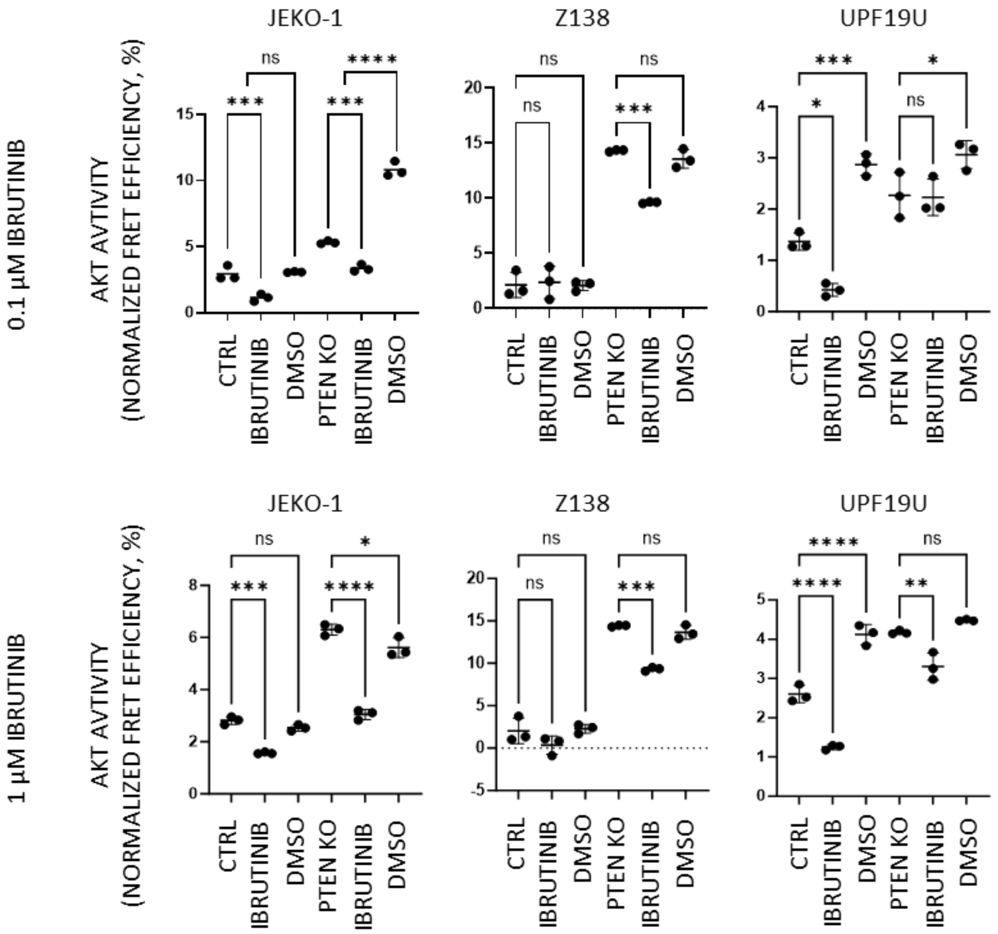


**Supplemental figure 4.** AKT activity is measured using genetically encoded FRET-based biosensor. AKT activity in *PTEN* KO cells is higher compared to respective unmodified cell lines. After exposure to BTK inhibitor ibrutinib for 3 hours (0.1 µM, 1 µM), AKT activity in *PTEN* KO cells remains higher than AKT activity in the respective CTRL cell lines; “ns” means “not significant”, * p<0.05, ** p<0.01, *** p<0.001, **** p<0.0001; N=3; (N) represents the number of biological replicates.

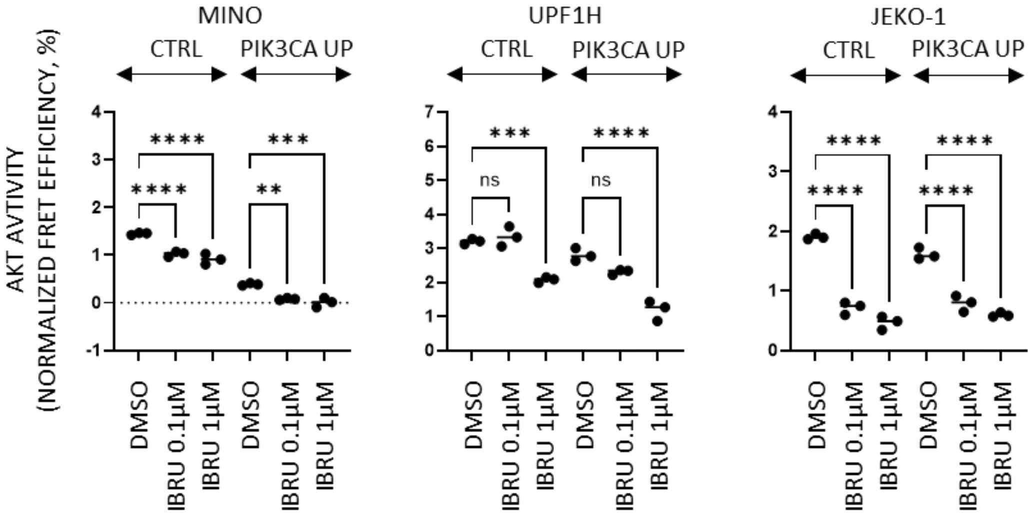


**Supplemental figure 5.** AKT activity is measured using genetically encoded FRET-based biosensor. Measurement of AKT activity after Ibrutinib (3 hours exposure) in 3 *PIK3CA* UP cell lines UPF1H, MINO and JEKO-1 by FRET assay showing that Ibrutinib decreases AKT activity in *PIK3CA* UP cell lines more than in CTRL in two of them; "ns" means "not significant", ** p<0.01, *** p<0.001, **** p<0.0001. N=3; (N) represents the number of biological replicates.

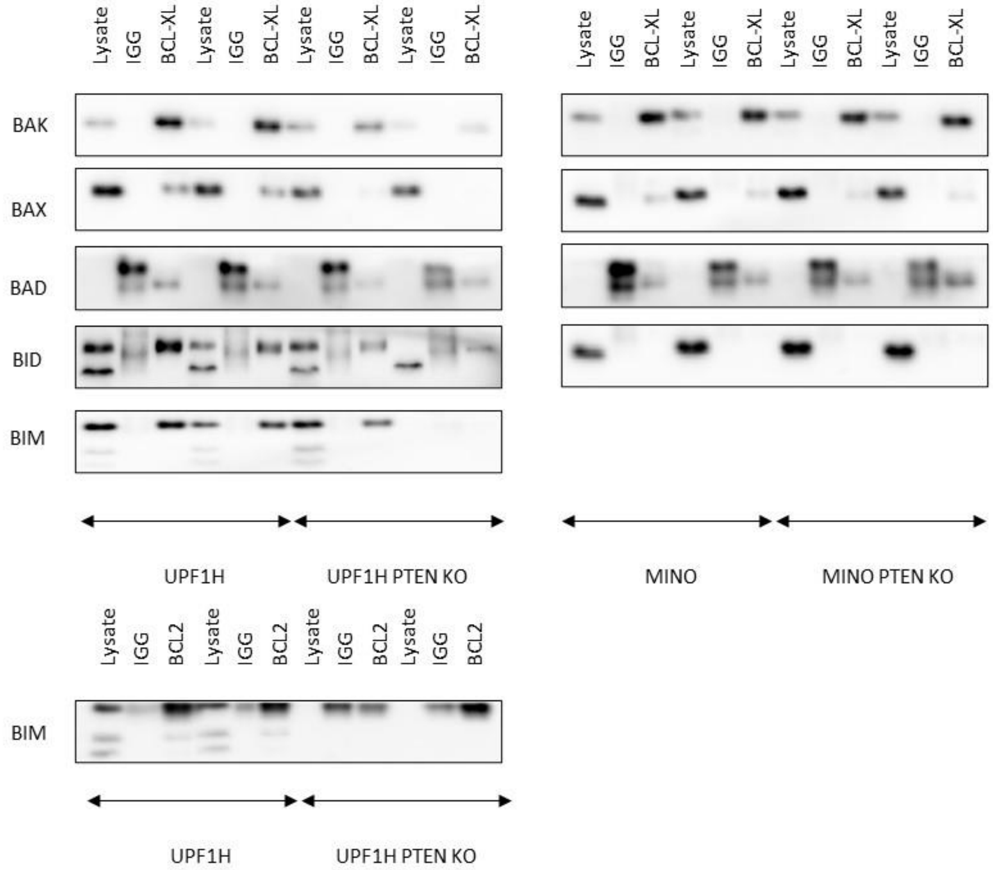

**Supplemental figure 6.** Co-immunoprecipitation of BCL-XL and BCL2. Decreased interaction of BIM, BAD, BID, BAK and BAX with BCL-XL in UPF1H *PTEN* KO. Decreased interaction of BAX with BCL-XL in MINO *PTEN* KO. Decreased interaction of BIM with BCL2 in UPF1H *PTEN* KO; for each sample N=2; (N) represents the number of biological replicates.

### SUPPLEMENTAL RFERENCES
